# Medial prefrontal cortex Notch1 signalling mediates methamphetamine-induced psychosis via Hes1-dependent suppression of GABA_B1_ receptor expression

**DOI:** 10.1101/2022.03.05.483073

**Authors:** Tong Ni, Li Zhu, Shuai Wang, Weili Zhu, Yanxue Xue, Yingjie Zhu, Dongliang Ma, Hongyan Wang, Fanglin Guan, Teng Chen

**Affiliations:** College of Forensic Medicine, Xi’an Jiaotong University Health Science Center, Xi’an, Shaanxi, 710061, PR China; The Key Laboratory of Health Ministry for Forensic Science, Xi’an Jiaotong University, Shaanxi, 710061, PR China; National Institute on Drug Dependence and Beijing Key Laboratory of Drug Dependence, Peking University, Beijing, 100191, PR China; Shenzhen Key Laboratory of Drug Addiction, CAS Key Laboratory of Brain Connectome and Manipulation, the Brain Cognition and Brain Disease Institute (BCBDI), Shenzhen Institutes of Advanced Technology, Chinese Academy of Sciences, Shenzhen, 518055, PR China; Shenzhen-Hong Kong Institute of Brain Science-Shenzhen Fundamental Research Institutions, Shenzhen, 518055, PR China; Programme in Neuroscience and Behavioral Disorders, Duke-NUS Medical School, 169857, Singapore; Department of Physiology, Yong Loo Lin School of Medicine, National University of Singapore, 117597, Singapore

**Keywords:** Methamphetamine, psychosis, Notch1, GABA_B1_ receptor, mPFC

## Abstract

Methamphetamine (METH), a widely abused stimulant drug, induces psychosis in approximately half of abusers; this effect is becoming a major concern for society. Although the Notch1 signalling pathway has been shown to play a part in the pathogenesis of some psychiatric disorders, its role in METH-induced psychosis (MIP) is still unknown. Here, we showed that Notch1 signalling was downregulated in the medial prefrontal cortex (mPFC) in a MIP mouse model. Direct genetic and pharmacological manipulations of Notch1 signalling bidirectionally altered MIP-related behaviours and governed neuronal activity in the mPFC. Moreover, Notch1 signalling negatively regulated GABAB1 receptor expression in the mPFC of MIP mice through Hes1, a transcriptional repressor in Notch1 signalling. Further, we show that Hes1 can directly bind to the GABA_B1_ receptor promoter. Notably, pharmacological regulation of the GABA_B_ receptor in the mPFC reversed the changes in MIP-related behaviours caused by the dysfunction of Notch1 signalling. Together, our findings uncover a previously unrecognised Notch1-Hes1-GABA_B1_ receptor-dependent mechanism involved in regulating mPFC neuronal activity and behavioural phenotypes in MIP. Our work provides mechanistic insight into the aetiology and pathophysiology of MIP.

## Introduction

Methamphetamine (METH) is the second most common illicit drug worldwide, with 33 million users^1^. Apart from its strong addictive properties, one of the most widely known health consequences associated with high-dose or chronic METH use is METH-induced psychosis (MIP), which affects between 26 and 46% of people with a METH dependence^2, 3^. MIP displays symptoms similar to schizophrenia (SCZ)^4^, including hyperactivity, agitation and cognitive deficiency, making it difficult to distinguish from SCZ clinically^5^. Despite the similarity between SCZ and MIP, the pathogenesis of MIP is different from SCZ and remains poorly understood.

Behavioural sensitization refers to the unique phenomenon whereby repeated exposure to a stimulus results in progressively increased behavioural activity in response to the stimulus following a period of abstinence^6^. Indeed, a single low-dose re-exposure to METH after decades of abstinence can cause MIP patients to display hyperactivity (manifested as hyperlocomotion) and a relapsed psychotic state^7, 8^. In animal studies, a METH-induced behavioural sensitization model also recapitulated MIP-related behaviours, including deficits in social behaviour^9^, cognitive functions^10^, and sensory gating^11^, which mimic the symptoms observed in patients with MIP and can be ameliorated by antipsychotic drugs^12^. Psychosis-related proteins were also detected in the medial prefrontal cortex (mPFC) of sensitized mice by proteomics studies^13, 14^. Hence, the METH-induced behavioural sensitization model is the most relevant model of MIP thus far^15^. Understanding the mechanism of METH sensitization is important for determining the aetiology of MIP that is distinct from SCZ.

The highly conserved Notch signal pathway is involved in multiple crucial processes, including stem cell fate determination and diversification during development^16^. After the Notch receptor (Notch1-4) interacts with one of its ligands (Delta or Jagged), the Notch intracellular domain is released by γ-secretase-mediated cleavage and moves into the nucleus, where it initiates transcription of Notch1 target genes, such as Hes1^17^. Notably, the Notch signalling, especially Notch1, could regulate synaptic plasticity and long-term memory in adult brain function from invertebrates to mammals^18, 19^. Interestingly, synaptic plasticity and memory are progressively affected in MIP^20, 21^, implying for the possible involvement of Notch1 in the neurological deficits associated with the disease. Moreover, Notch1 imbalances have been evidenced in patients and animal models affected by psychoses, such as SCZ^22, 23^, depression and anxiety^24^. However, whether and how the Notch1 pathway is involved in MIP remains unclear.

It has been proposed that damage of cortical GABAergic function leads to dysregulation and imbalance of glutamatergic and GABAergic cortical signals, resulting in MIP^2^. Previous studies identified GABA transporters and receptors expression changes in the mPFC^25^, nucleus accumbens (NAc)^26^ and hippocampus (Hip)^27^ following METH sensitization in mice. In addition, researchers have found an association between Notch1 signalling and GABA transporters or receptors^28, 29^. Therefore, we hypothesize that Notch1 signalling is involved in MIP by regulating GABAergic genes expression. In this study, we investigated the neuron-specific changes in Notch1 expression on MIP-associated behaviours and elucidate the mechanism by which the Notch1 signalling pathway regulates MIP via the GABAergic system. Our findings provide insights into the pathogenesis of MIP and may facilitate the development of improved treatments.

## Materials and methods

### Animals

Male C57BL/6J mice (2 months old and weighing 20-25 g) were purchased from Beijing Vital River Laboratory Animal Technology Co., Ltd. (Beijing, China). Animals were housed on a light/dark cycle of 12 h/12 h in standard group cages (≤ 5 mice/cage) and had unrestricted access to food and water. Mice were randomly assigned to different experimental groups. All the animal protocols used in this study were approved by the Institutional Animal Care and Use Committee of Xi’an Jiaotong University and followed the guidelines established by the National Institutes of Health.

### Drug treatment

For METH sensitization, METH was purchased from the National Institute for Control of Pharmaceutical and Biological Products (Beijing, China) and dissolved in 0.9% NaCl (saline). METH (1 mg/kg or 5 mg/kg, dissolved in saline) and saline were each administered via intraperitoneal (i.p.) injection. For the SCZ animal model, we used NMDA receptor antagonist MK-801 (Abcam, USA, 1 mg/kg, i.p.) once daily^30^ for 21 consecutive days. For the microinjection, DAPT (Selleck, USA), a key enzyme inhibitor of Notch signal pathway, is dissolved in 90% DMSO (sigma, USA) prepared in 0.1M sterile phosphate-buffered saline (PBS)^31^ at 30 μg/μL. DAPT or vehicle was administered via bilateral intracranial microinjections at 0.5 μL/hemisphere. The GABA_B_ receptor agonist baclofen (Sigma, USA) and the GABA_B_ receptor antagonist phaclofen (Sigma, USA) were dissolved in saline and administered at doses of 0.06 nmol/0.2 μL/mouse^32^ and 0.1nmol /0.2μL/ mouse^33^, respectively.

### Cannulation and microinjection

Briefly, the mice were anaesthetized using 1.5% isoflurane and placed in an automated stereotaxic instrument (RWD Life Science, China). A 0.8-cm-long stainless steel cannula was unilaterally implanted in the mPFC [anteroposterior (AP): + 2.00 mm; mediolateral (ML): ± 0.50 mm; dorsoventral (DV): - 2.4 mm]^34^. After 1 week of recovery, DAPT or vehicle and baclofen, phaclofen or saline were infused into the mPFC 30 min before 1 mg/kg METH (or saline) was injected intraperitoneally. For the intracerebral infusions, the solutions were injected at a rate of 0.1 μL/minute. The injection cannula was left in place for an additional 5 minutes to minimise the efflux of the drug.

### Adeno-associated virus generation and injection

We used AAV2/8 expressing synapsin promoter with Notch1 intracellular domain (NICD) to overexpress NICD in neurons (syn-NICD-OE), shRNA1 (GCCTCAATATTCCTTACAA) to knock down the NICD expression in neurons (syn-NICD-shRNA) and shRNA2 (TGAAAGTCTAAGCCAACTGAA) to knock down the expression of Hes1 in neurons (syn-Hes1-shRNA). The same vector backbone was used to generate a negative control. Mice were anaesthetized with 1.5% isoflurane before AAV injection. AAVs were injected bilaterally into the mPFC at the corresponding coordinates (AP: +2.05 mm; ML: ±0.27 mm; DV: −2.10 mm)^35^. The AAV vectors (200 nL) were infused slowly per side over 4’min into the targets using a micro-infusion pump with a 10 μL Hamilton syringe. The microsyringe was left in place for 6 min to allow the viral vectors to diffuse after microinjection. Behavioural testing was initiated four weeks after injections.

### Behavioural Testing

METH-induced sensitization was performed as previously described (Fig. 1A)^25^. Briefly, METH-sensitized group received repeated doses of METH on days 1 and 7; 5 mg/kg, i.p. on days 2–6. The acute METH and saline groups were injected with saline for 7 days. Then, after a 2-week withdrawal period, the METH-sensitized and acute METH group received the same challenge dose of METH (1 mg/kg), while the saline group received one injection of saline on day 23 (Fig. 1A). Horizontal locomotor activities were recorded in metal test chambers (43 cm × 43 cm × 43 cm) and analysed for 60 minutes after 1 mg/kg METH injections using a smart 2.5 video tracking system. The amount of time travelled in the centre zone (21.50 cm×21.50 cm) are interpreted as measures of anxiety-like behavior^36^.

**Figure 1.**
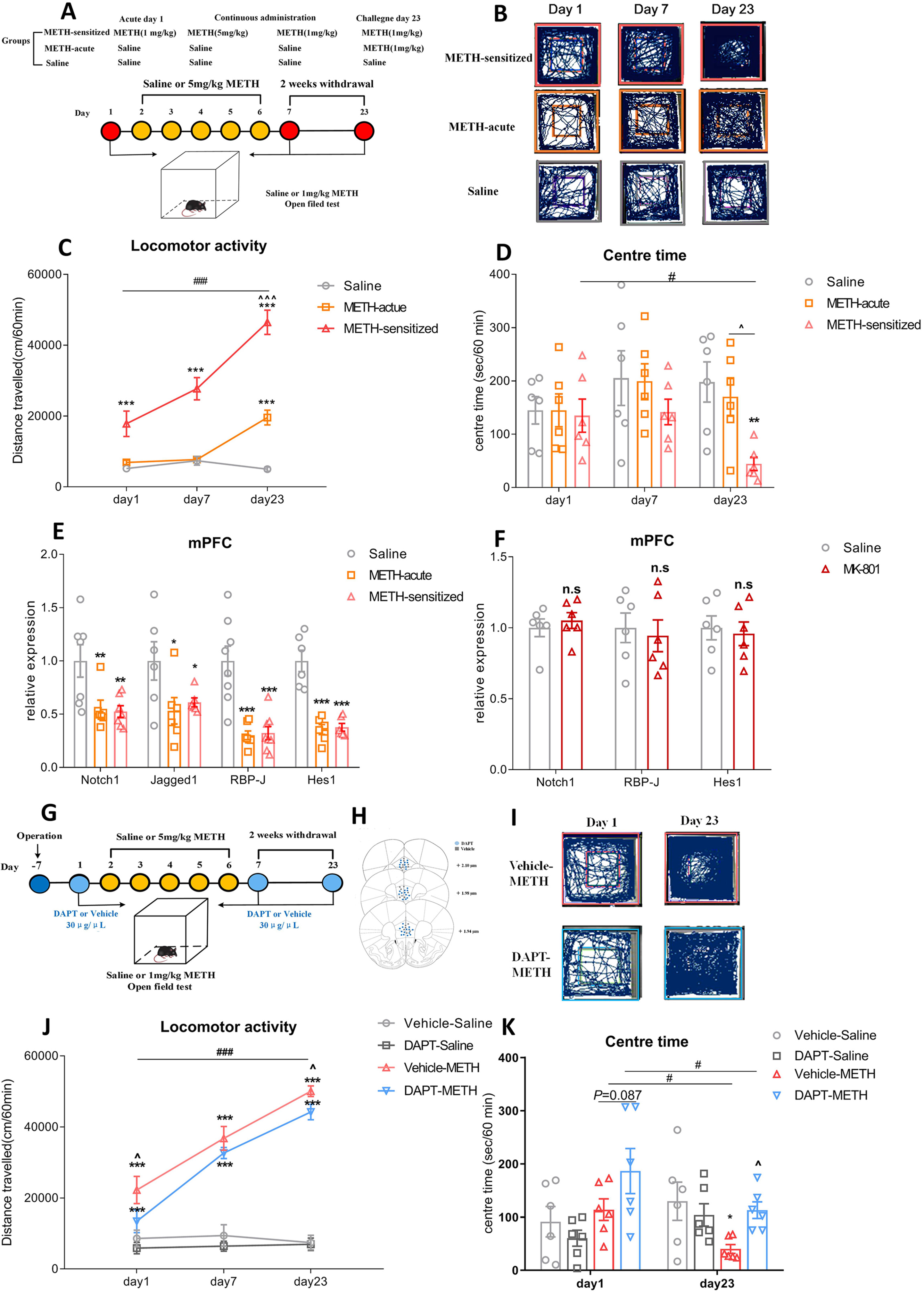
The involvement of the Notch1 signalling pathway in the mPFC of METH-induced sensitized mice. 1A-1E. Changes in Notch1 signalling in the mPFC of METH-sensitized mice. **1A.** Procedure for generating the METH-induced locomotor sensitization model. **1B.** Representative mouse tracks from days 1, 7 and 23 were illustrated. **1C.** METH-induced locomotor sensitization in mice. One-way repeated-measures ANOVA revealed a significant main effect of METH [F _(2, 21)_ = 47.41, *P <* 0.001]. Subsequent post hoc LSD comparisons found significantly greater locomotor activity in the METH-sensitized group than in the saline group on day 1, day 7 and day 23. The METH-sensitized group showed a significant increase in locomotor activity compared with the METH-acute group on day 23 and itself on day 1. **1D.** The amount of time mice spent in the centre zone significantly decreased in the METH-sensitized group compared with the saline and acute METH group on day 23 [F _(2, 15)_ = 3.66, *P <* 0.05]. **1E.** Changes in the expression of Notch1 signalling pathway components following METH sensitization in the mPFC. There was significant downregulation of Notch1 [F _(2, 17)_ = 6.41, *P <* 0.01], Jagged1 [F _(2, 15)_ = 3.81, *P <* 0.05], RBP-J [F _(2, 20)_ = 18.05, *P <* 0.001], and Hes1 [F _(2, 15)_ = 29.68, *P <* 0.001] in both the METH-acute and METH-sensitized groups by one-way ANOVA**. 1F.** Changes in the expression of Notch1 signalling components in the mPFC of the MK-801-induced SCZ animal model. There were no significant changes (n.s.) of Notch1 (t_10_ = −0.62), RBP-J (t_10_ = 0.37) or Hes1 (t_10_ = 0.36) in the mPFC of MK-801-treated mice. **1G-1K.** Intra-mPFC DAPT treatment attenuated METH-induced locomotor sensitization. **1G.** Procedure for the administration of DAPT in the mPFC in METH-induced sensitization mice. **1H.** Location of the DAPT and vehicle microinjection cannula tips in the mPFC. **1I.** Representative tracks of vehicle-METH and DAPT-METH mice on day 1 and day 23. **1J.** Intra-mPFC infusion of DAPT significantly attenuated the hyperlocomotion evoked by METH on day 1 and day 23. Mixed-design ANOVA with a LSD post hoc multiple comparison revealed significant main effects of DAPT [F _(1, 21)_ = 5.38, *P <* 0.05]; METH [F _(1, 21)_ = 206.21, *P <* 0.001]; but not DAPT×METH [F _(1, 21)_ = 1.36, *P >* 0.05]. **1K.** The time mice spent in the centre zone was higher in the DAPT-METH group than in the vehicle-METH group on day 23. Mixed-design ANOVA with LSD post hoc multiple comparisons show the main effect of DAPT [F _(1, 20)_ = 1.24, *P >* 0.05]; METH [F _(1, 20)_ = 0.74, *P >* 0.05] and DAPT×METH [F _(1, 20)_ = 6.45, *P <* 0.05]. **\****P <* 0.05, \*\**P <* 0.01, \*\*\**P <* 0.001 vs. paired saline group; ∧*P <* 0.05 vs. paired METH group; #*P <* 0.05, ###*P <* 0.001 vs. same group on day 1. Data were presented as mean ± S.E.M, n = 6.

The specific methods of other behavioural testing, such as open filed test, novel object recognition (NOR), Y maze test, elevated plus maze (EPM), tail suspension test (TST) and forced swimming test (FST), were seen in the supplement methods.

### Quantitative real-time reverse transcription PCR (RT-qPCR)

Mice were sacrificed 1 hour after the last METH/saline injection, and the brains were rapidly removed. The mPFC, NAc and Hip were dissected out based on mouse brain structure and landmarks under dissecting microscopy. Total RNA was isolated using a DNA/RNA/protein kit (Omega, USA). Total RNA was reverse transcribed into 10 μL of complementary DNA (cDNA) with PrimeScript^TM^ RT Master Mix (Takara Biomedical Technology, China) at 37°C for 15 min, 85°C for 5 s, and 4°C for 5 min. RT-qPCR for mRNA detection was performed with the SYBR Premix Ex Taq II (Takara Biomedical Technology, China) using Bio-Rad iQ5 detection instrument (Bio-Rad, USA) under the following conditions: 95°C for 30 s, 40 cycles of 95°C for 15 s, 60°C for 30 s and 72°C for 30 s. Gene expression was analysed as previously described^37^. The sequences of the primer pairs were shown in supplement Table 1.

### Western blot

Total mPFC tissue was collected 1 hour after the last injection, and the protein concentration in each sample was determined using a BCA protein assay kit (Pierce, Rockland, IL, USA). These protein samples were separated using 10% SDS–PAGE and transferred to nitrocellulose membranes (Millipore, USA). Then, the membranes were blocked for 3 h in 5% (w/v) milk at room temperature and incubated overnight at 4°C with the following primary antibodies: rabbit anti-Notch1 (1:1000, Abcam, ab8925), rabbit anti-Hes1 (1:1000, Abcam, ab71559), and mouse anti-GABA_B1_ receptor (1:2000, Abcam, ab55051). Membranes were then washed with TBST and probed with the appropriate horseradish peroxidase-conjugated secondary antibodies (1:2000) for 1.5 h at room temperature. Proteins were detected with an enhanced chemiluminescence assay kit (ECL Plus, Millipore Corporation, USA). Signals were visualized using ImageLab 1.46 (BioRad, USA).

### Chromatin immunoprecipitation (ChIP)-qPCR

ChIP assays were performed using a ChIP assay kit (Millipore EZ-CHIP 17-371, USA) according to the manufacturer’s instructions^38^. Briefly, crosslinking was performed with 1% formalin, the mPFC was lysed in SDS buffer, and sonication was used to fragment the DNA to an average length of 200 to 500 base pairs. The input group accounted for 1% of the total DNA, while IgG and Hes1 (1:10, Cell Signaling Technology, #11988, USA) antibodies were added as negative controls and samples. After purification, RT–qPCR was performed to detect the protein binding sites of the DNA samples. The ChIP signal was calculated as follows: % input = 1% × 2 ∧ (CT_input_ - CT_sample_). The sequences of the RT-PCR primers used for GABA_B1_ receptor promoter were shown in supplement Table 1.

### Fiber photometry

Mice were unilaterally injected with 0.3 μL of AAV-syn-GCaMP7f and 0.2 μL of AAV-syn-NICD-shRNA-mCherry or control into the ipsilateral mPFC. A unilateral optical fiber (200 μm core, 0.39 numerical aperture (NA), RWD, China) was implanted at these coordinates and secured in place using dental cement after two weeks of recovery from AAV-micro-injection. Then, the mice recovered in their home cage for 7 days before beginning behavioural testing. A fiber photometry system (R810, RWD Life Science, China) was used to record the fluorescence signal (GCaMP7f), which was produced by an exciting laser beam from 470 nm LED light and 410 nm LED light^39^. On the experimental day, mice were allowed to acclimate in the behavioural testing chamber for 30 min. After the acclimation period, baseline fluorescence was recorded for 5 minutes. Then, the mice were injected with METH (1 mg/kg, i.p.). Fluorescence was then recorded with the optical fiber for 15 min after the administration of METH. ΔF/F was calculated according to (470 nm signal-fitted 410 nm signal)/ (fitted 410 nm signal). The standard Z score calculation method was performed in MATLAB 2014. The formula was as follows: Z score = (x-mean)/std, x = ΔF/F.

### Statistical analysis

Statistical analyses were performed using SPSS (version 18.0). One-way or mixed-designed repeated ANOVA with multiple comparisons followed by LSD post hoc were applied to the METH-induced locomotor sensitization data. Other data from behavioural tests, RT–qPCR and Western blot were analysed by one-way or two-way ANOVA followed by LSD post hoc test. The averaged Z scores of fiber photometry were compared by a paired-sample t test, and ChIP-qPCR data were analysed by student’s t test. All data were expressed as the mean ± standard error of the mean (SEM), and *P <* 0.05 was considered statistically significant.

## Results

### The Notch1 signalling pathway was downregulated in the mPFC of MIP mice

Here, mice were treated with METH in a standard protocol to examine METH-induced locomotor sensitization (Fig. 1A), which was considered as a MIP model^25^. Repeated intermittent METH administration led to a progressive augmentation of behavioural changes compared with the saline group, and mice showed a significant sensitization response compared with the acute METH group on day 23 (Fig. 1B-C). Meanwhile, the time that mice spent in the centre area of the open field was significantly decreased following METH challenge (Fig. 1D), which indicated that METH challenge induced increased anxiety-like behaviour in mice. Moreover, the SCZ animal model induced by 21-day consecutive treatment of MK-801^30^ also exhibited the hyperlocomotion and decreased centre time (Fig. S1A).

We next examined the mRNA levels of Notch1 signalling in the mPFC, NAc, and Hip, which are psychosis-related brain regions. Our results showed that both acute METH treatment and METH sensitization induced a significant downregulation of Notch1, Jagged1, RBP-J and Hes1 in the mPFC (Fig. 1E), but not all of these genes showed significant changes in the NAc or Hip (Fig. S1B-C). Therefore, we chose the mPFC as the target region to further examine the role of the Notch1 pathway in MIP. We also measured the changes in expression of Notch1 signalling in the mPFC of mice with MK-801. Surprisingly, there were no significant differences of Notch1 signalling in the mPFC after MK-801 treatment (Fig. 1F). Similarly, the data from PsychENCODE Consortium in the NIMH Repository (http://psychencode.org), which collected the transcription profiles of prefrontal cortex tissue from postmortem SCZ patients^40^, showed no difference in Notch1 signalling between SCZ patients and healthy controls (Fig. S1D). These results suggested that specific alterations in Notch1 signalling at the mPFC in the MIP mouse model.

### Differential expression of Notch1 signalling in mPFC was capable of regulating MIP-related behaviours

#### Inhibition of Notch1 signalling in the mPFC attenuated METH-induced locomotor sensitization

To test whether inhibition of Notch1 signalling in mPFC could affect METH-induced locomotor sensitization, we first administered the Notch signalling inhibitor DAPT^31^. Vehicle or DAPT (30 μg/μL) was administered to the mPFC 30 min before saline or 1 mg/kg METH treatment (Fig. 1G-H). DAPT itself did not induce any locomotion responses in mice (Fig. 1J). However, DAPT suppressed METH-induced hyperlocomotion on acute and challenge days (Fig. 1I-J) and enhanced the time spent in the centre area in response to METH on challenge day (Fig. 1K).

We used syn-NICD-shRNA to inhibit the neuronal expression of Notch1 intracellular domain (NICD) in the mPFC (Fig. 2A) and obtained similar results as that of DAPT treatment. The mRNA and protein levels of NICD were significantly decreased in the syn-NICD-shRNA group (Fig. 2B). Both groups were then subjected to repeated intermittent METH treatment, and the locomotor activity of the mice was tested (Fig. 2C-D). Downregulation of NICD in the mPFC significantly reduced the total distance travelled in response to METH on the acute and challenge day (Fig. 2C-D) and increased the time spent in the centre area (Fig. 2E).

**Figure 2.**
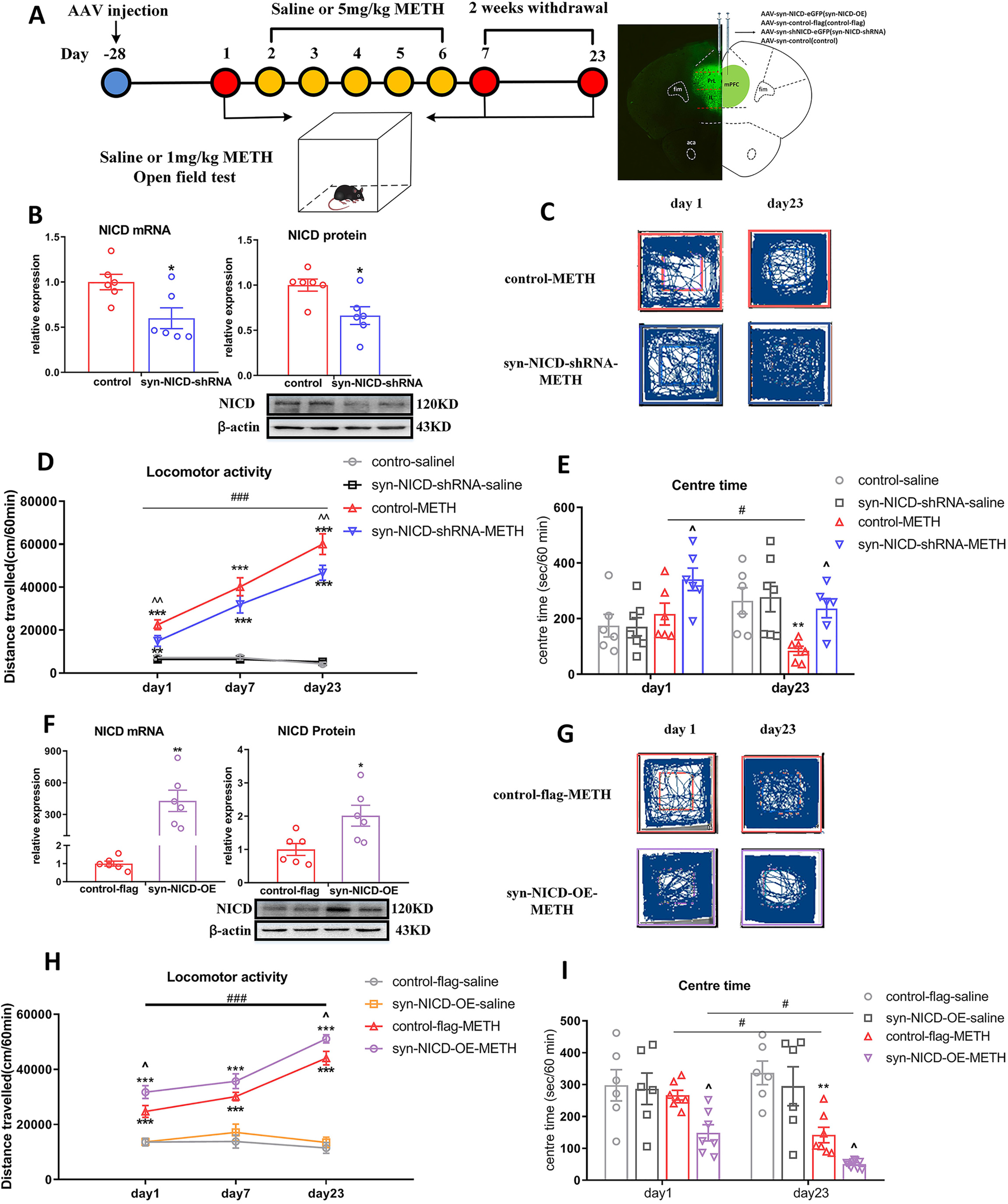
Manipulating Notch1 expression affects METH-induced locomotor sensitization. 2A. Timeline of AAV injection, behavioural tests and location of AAV expression. **2B-2E.** Inhibition of NICD in the mPFC suppressed METH-induced locomotor activity. **2B.** NICD mRNA (t_10_ = 2.79, \**P <* 0.05) and protein (t_10_ = 2.84, \**P <* 0.05) levels were significantly lower in syn-NICD-shRNA group as indicated by student’s t test. **2C, 2G**. Representative paths of mice in the syn-NICD-shRNA-METH and control-METH or syn-NICD-OE-METH and control-flag-METH groups on day 1 and day 23. **2D.** Downregulation of NICD in mPFC neurons significantly decreased locomotion on day 1 and day 23. Mixed-design ANOVA with a LSD post hoc multiple comparison revealed significant main effects of AAV [F _(1, 20)_ = 8.95, *P <* 0.01]; METH [F _(1, 20)_ = 323.81, *P <* 0.001]; and AAV×METH [F _(1, 20)_ = 8.10, *P <* 0.05]. **2E.** Time spent in the centre zone was significantly higher in the syn-NICD-shRNA-METH group than in the control-METH group on day 1 and day 23 [main effect of AAV, F _(1, 21)_ = 7.61, *P <* 0.05; METH, F _(1, 21)_ = 0.01, *P >* 0.05; AAV×METH, F _(1, 21)_ = 7.18, *P <* 0.05]. **2F-2I**. Overexpression of Notch1 in the mPFC enhanced METH-induced locomotor activity. **2F.** NICD mRNA (t_10_ = −4.25, \*\**P <* 0.01) and protein (t_10_ = −2.81, \**P <* 0.05) levels were significantly higher in the mPFC of the syn-NICD-OE group than the control-flag group as determined by student’s t test. **2H.** Overexpression of NICD significantly increased locomotion on day 1 and day 23 compared with the control-flag-METH group. Mixed-design ANOVA with a LSD post hoc multiple comparison revealed significant main effects of AAV [F _(1, 20)_ = 8.28, *P <* 0.01]; METH [F _(1, 20)_ = 241.37, *P <* 0.001]; but not the AAV×METH [F _(1, 20)_ = 2.70, *P >* 0.05]. **2I.** The time that mice spent in the centre zone was significantly downregulated in the syn-NICD-OE-METH group compared with the control-flag-METH group on day 1 and day 23 [main effect of AAV, F _(1, 22)_ = 6.50, *P <* 0.05; METH, F _(1, 22)_ = 32.14, *P <* 0.001; AAV×METH, F _(1, 22)_ = 2.09, *P >* 0.05]. \**P <* 0.05, \*\**P <* 0.01, \*\*\**P <* 0.001 vs. paired saline group; ∧*P* < 0.05 vs. paired METH group; #*P <* 0.05, ###*P <* 0.001 vs. same group on day 1. Data were presented as mean ± S.E.M, n = 6-8.

#### Overexpression of Notch1 signalling in the mPFC enhanced METH-induced locomotor sensitization

We further tested whether overexpressing NICD (syn-NICD-OE) in mPFC neurons could enhance METH-induced locomotor sensitization. Both the mRNA and protein levels of NICD in the mPFC were significantly upregulated after AAV microinjection (Fig. 2F). As expected, the overexpression of NICD in the mPFC significantly enhanced the locomotor distance on acute and challenge day (Fig. 2G-H) and decreased the time spent in the centre area compared to the control METH group (Fig. 2I).

#### Regulation of Notch1 signalling expression in the mPFC influenced other MIP-related behaviours

MIP also exhibited other behaviours relevant to psychiatric symptoms, such as cognitive impairment, depression and anxiety^41^. Thus, we examined the influences of Notch1 signalling on these behaviours of mice after METH treatment (Fig. S2A). The syn-NICD-shRNA group attenuated the MIP-related cognitive impairment of mice in both NOR (Fig. S2C) and Y maze test (Fig. S2D). Moreover, the reduction of NICD in mPFC also reversed the MIP-related anxiety-like behaviours in the EPM (Fig. S3A-B) and depression-like behaviours in the TST (Fig. S3C) and FST (Fig. S3D) respectively. In contrast, syn-NICD-OE aggravated MIP-related behaviours in the NOR (Fig. S2F), Y maze (Fig. S2G), EPM (Fig. S3E-F), TST and FST (Fig. S3G-H). All these data corroborate the idea that differential expression of Notch1 signalling in the mPFC was capable of regulating MIP.

### Downregulation of Notch1 signalling in the mPFC attenuated neuronal activity in MIP

The dysregulation of mPFC neuronal activity is a key factor causing MIP^42^. Therefore, we characterised whether mPFC neuronal activity was modulated by regulating Notch1 signalling in MIP (Fig. 3A). We co-injected the AAV-syn-GCaMP7f and syn-NICD-shRNA-mCherry into the mPFC region of mice, nearly 95% of the GCaMP7-positive cells also expressed mCherry (Fig. 3B). We further employed fiber photometry to monitor the Ca^2+^ signals of mPFC neurons in the acute phase and on the challenge day. Syn-NICD-shRNA induced a significant reduction of mPFC neurons activity in response to acute METH exposure (Fig. 3C, 3E, 3G). On the challenge day, the fluorescent signal was first reduced within approximately five minutes and then increased gradually (Fig. 3D, 3F). However, neuronal activity tested in the syn-NICD-shRNA mice showed no increasing trend after METH injection on day 23 (Fig. 3D, 3F) and demonstrated a significant reduction compared to what was before METH treatment (Fig. 3H). These data identified that the downregulation of NICD in the mPFC could directly attenuate neuronal activity in mice with MIP.

**Figure 3.**
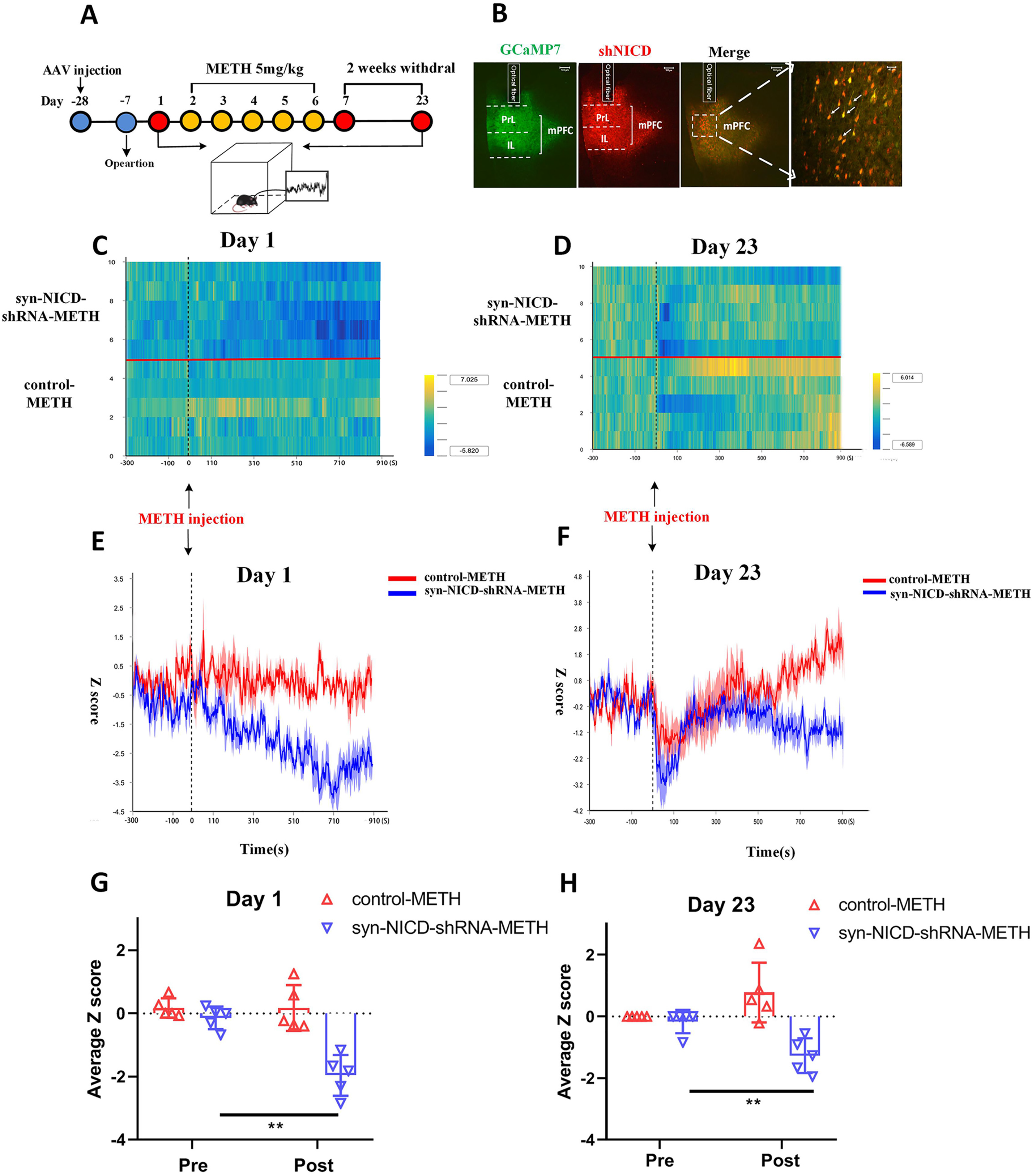
Inhibition of the Notch1 signalling attenuates mPFC neuronal activity of METH-induced sensitized mice. 3A. Experimental design for recording GCaMP activity from mPFC neurons which reduced the Notch1 signalling during METH sensitization. Calcium-dependent (470 nm) and calcium-independent (410 nm) fluorescence signals were recorded before and after administration of METH (1 mg/kg) in the open field. **3B.** Fluorescence images in the mPFC from mice that received unilateral infusion of syn-GCaMP7f (green) and syn-NICD-shRNA (red) virus, scale bar = 100 µm. Arrows indicate neurons both expressed GCaMP7f and NICD-shRNA signals, scale bar = 40 µm. **3C-3D.** Heatmap illustration of Ca^2+^ signals aligned to the initiation of trials on day 1 and day 23. Each row plots one trial, and a total of 10 trials were illustrated. The colour scale on the right indicates Z scores. **3E-3F.** Average traces of calcium signals from control mPFC neurons (red line) and inhibitory NICD mPFC neurons (blue line) were recorded before (5 minutes) and after METH treatment (15 minutes) on days 1 and 23. **3G-3H.** The average Z scores from (E) and (F) in the open field before (pre) and after (post) METH injection on day 1 and day 23. The fiber photometry signal of mPFC neurons in the syn-NICD-shRNA-METH group was significantly reduced compared with the baseline signal on day 1 and day 23, according to a paired-sample t test (t _day1_ = 7.953, t _day23_ = 4.58, \*\**P <* 0.01 vs. before METH injection). Traces represent mean ± SEM (3E, 3F). Error bars represent mean ± SEM (3G, 3H), n = 5.

### The Notch1 signalling pathway in the mPFC negatively regulated GABA_B1_ receptor expression

Previously, differential expression of GABA transporters (GAT1 and GAT3), ionotropic GABA_A_ receptor subunits (α3 and β1), and metabotropic GABA_B_ receptors were found in the mPFC of the same animal model of MIP^43^. Moreover, Notch1 signalling was associated with GABA receptors and transporters expression^28, 29, 44^. We speculated that the GABAergic system may underlie the mechanism through which Notch1 signalling regulated MIP. Indeed, we found that the expressions of GABA receptors and transporters in the mPFC decreased in both the acute METH and METH-sensitized groups (Fig. 4A). GABA_Aβ1_, GABA_B1_ and GAT1 were significantly upregulated in the METH-sensitized group compared to the acute METH group (Fig. 4A).

**Figure 4.**
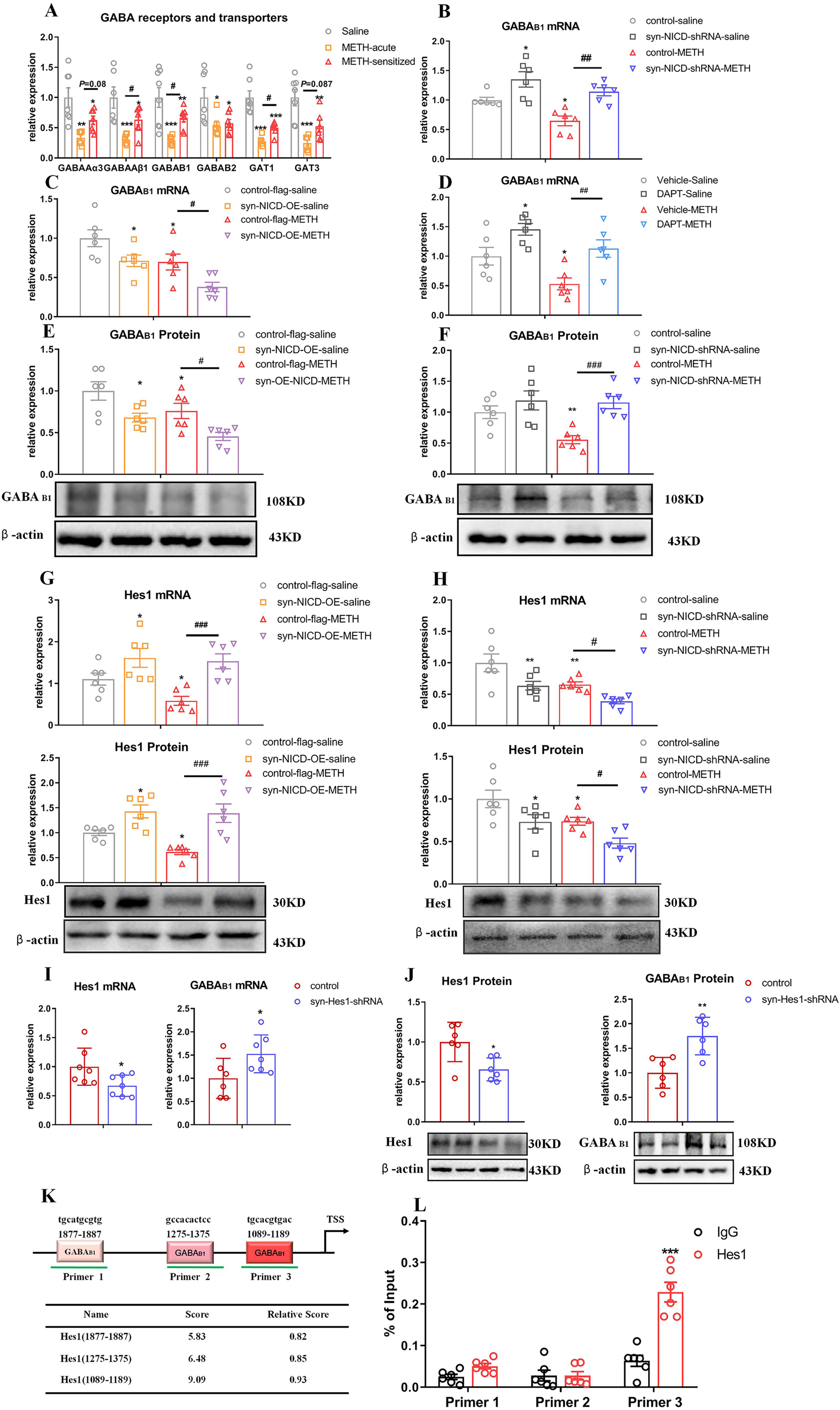
The Notch1 signalling pathway regulated the GABA_B1_ receptor via hes1 in the mPFC of METH-induced sensitized mice. 4A. GABA receptors and transporters expression changes following METH sensitization in the mPFC. One-way ANOVA showed the significant downregulation effects of METH on GABA_Aα3_ [F _(2, 17)_ = 9.17, *P <* 0.05], GABA_Aβ1_ [F _(2, 17)_ = 10.38, *P <* 0.05], GABA_B1_ [F_(2, 18)_ = 11.66, *P <* 0.05], GABA_B2_ [F _(2, 17)_ = 5.36, *P <* 0.05], GAT1 [F _(2, 17)_ = 25.79, *P* < 0.05] and GAT3 [F _(2, 16)_ = 13.48, *P <* 0.05]. Subsequent post hoc LSD comparisons found higher expression changes of GABA_A_β1, GABA_B1_ and GAT1 between the METH-sensitized group and acute METH group. \**P <* 0.05, \*\**P <* 0.01. \*\*\**P* < 0.001 vs. saline group. #*P <* 0.05 METH-sensitized group vs. acute METH group. **4B-4F.** GABA_B1_ receptor expression changes following regulation of the Notch1 signalling pathway in the mPFC of MIP mice. **4B, 4F.** Two-way ANOVA followed by LSD test showed significant upregulation of the GABA_B1_ receptor in the syn-NICD-shRNA group at both the mRNA and protein levels in the mPFC of MIP mice [4B. main effect of AAV, F _(1, 20)_ = 23.29, *P <* 0.001; METH, F _(1, 20)_ = 10.27, *P <* 0.01; AAV×METH, *P >* 0.05] and protein level [4F. main effect of AAV, F _(1, 20)_ = 13.09, *P <* 0.01; METH, F _(1, 20)_ = 4.78, *P <* 0.05; AAV×METH, *P >* 0.05]. **4C, 4E.** There was significant downregulation of the GABA_B1_ receptor at both the mRNA [4C. main effect of AAV, F _(1, 20)_ = 12.03, *P <* 0.01; METH, F _(1, 20)_ = 13.28, *P <* 0.01; AAV×METH, *P >* 0.05] and protein [4E. main effect of AAV, F _(1, 20)_ = 15.31, *P <* 0.01; METH, F _(1, 20)_ = 8.44, *P <* 0.01; AAV×METH, *P >* 0.05] levels in the mPFC of MIP mice in the syn-NICD-OE group. **4D.** DAPT in the mPFC induced increased GABA_B1_ receptor mRNA levels in the mPFC of MIP mice [main effect of DAPT, F _(1, 20)_ = 17.38, *P <* 0.01; main effect of METH, F _(1, 20)_ = 9.88, *P <* 0.01; DAPT×METH, *P >* 0.05]. **4G-4H.** Hes1 expression changes following regulation of the Notch1 signalling pathway in the mPFC of MIP mice. **4G.** Hes1 mRNA [main effect of AAV, F _(1, 20)_ = 17.47, *P <* 0.001; METH, F _(1, 20)_ = 2.89*, P >* 0.05; AAV×METH, *P >* 0.05] and protein [main effect of AAV, F _(1, 20)_ = 25.73, *P <* 0.05; METH, F _(1, 20)_ = 3.16, *P* > 0.05; AAV×METH, *P >* 0.05] levels were increased in the syn-NICD-OE group of mice. **4H.** Hes1 mRNA [main effect of AAV, F _(1, 20)_ = 14.17, *P <* 0.01; METH, F_(1, 20)_ = 12.56, *P <* 0.01; AAV×METH, *P >* 0.05] and protein [main effect of AAV, F _(1, 20)_ = 11.84, *P <* 0.01; METH, F _(1, 20)_ = 11.27, *P <* 0.01; AAV×METH, *P >* 0.05] levels were decreased in the syn-NICD-shRNA group of mice. **4I-4J.** Hes1 mRNA (t_12_ = 2.35, \**P* < 0.05) and protein (t_10_ = 2.95, \**P <* 0.05) levels were significantly lower in syn-Hes1-shRNA mice than control mice by student’s t test. GABA_B1_ receptor mRNA (t_11_ = −2.27, \**P <* 0.05) and protein (t_10_ = −3.71, \*\**P <* 0.01) levels were increased in the syn-Hes1-shRNA group. **4K-4L.** The results of ChIP-qPCR assays indicated that Hes1 bound to the promoter of the GABA_B1_ receptor. **4K.** Schematic diagram and table of the GABA_B1_ promoter region showing potential binding sites of Hes1 predicted by the JASPAR database. **4L.** The naive mice mPFC were subjected to ChIP assay. Three paired primers were designed near the predicted binding sites of Hes1. ChIP-qPCR assays verified the association of Hes1 and the promoter of the GABA_B1_ gene (Primer 3, t_10_ = −6.11, \*\*\**P <* 0.001). \**P <* 0.05, \*\**P <* 0.01, \*\*\**P <* 0.001 vs. control saline group. #*P <* 0.05, ##*P <* 0.01 vs. paired METH group. The results are expressed as the mean ± SEM, n = 6-7.

We then assessed whether the changes in the expression of GABA receptors and transporters were regulated by Notch1 signalling in the mPFC. We found that syn-NICD-shRNA in the mPFC significantly reversed the reduction in GABA_Aβ1_, GABA_Aα3_ and GABA_B1_ in response to METH (Fig. 4B, Fig. S4A-E), while syn-NICD-OE only significantly decreased GABA_B1_ expression (Fig. 4C, Fig. S4F-J). In addition, the direct injection of DAPT into the mPFC showed the same effects as syn-NICD-shRNA on GABAergic genes (Fig. 4D, Fig. S4K-O). Western blotting confirmed the changes in the expression of the GABA_B1_ receptor when NICD expression in the mPFC as manipulated (Fig. 4E-F). As expected, syn-NICD-OE reduced the expression level of the GABA_B1_ receptor in the mPFC (Fig. 4E), whereas syn-NICD-shRNA group showed the increased level of GABA_B1_ receptor compared with the control group (Fig. 4F). Taken together, the evidence suggested that the regulation of Notch1 signalling in the mPFC negatively regulated the expression of the GABA_B1_ receptor.

### Notch1 signalling regulated the expression of the GABA_B1_ receptor directly via Hes1

Hes1, a downstream transcriptional repressor of Notch1 signalling, expressed in neurons and controls GABAergic differentiation^45^. Therefore, we surmised that the transcriptional repressor Hes1 might be the key factor negatively regulating GABA_B1_ receptor expression. To test this hypothesis, we measured the mRNA and protein levels of Hes1 in the mPFC. Remarkably, Hes1 was upregulated in the syn-NICD-OE group, in which the abundance of GABA_B1_ receptors was decreased (Fig. 4G). In contrast, Hes1 was reduced in the syn-NICD-shRNA group with higher levels of GABA_B1_ receptors (Fig. 4H). Furthermore, shRNA was used to suppress Hes1 expression in the mPFC (Fig. 4I-J) and GABA_B1_ receptor expression was significantly increased in the syn-Hes1-shRNA group (Fig. 4I-J).

The JASPAR database predicted 3 potential binding sites for Hes1 on the GABA_B1_ promoter (Fig. 4K). We performed ChIP-qPCR to determine whether Hes1 directly bind to GABA_B1_ receptor promoter to regulate expression. According to the ChIP-qPCR results, there was significant enrichment of Hes1 at the specific binding sites (primer 3) of the GABA_B1_ receptor promoter (Fig. 4L). Taken together, these results suggest that Hes1 transcriptionally inhibited the expression of the GABA_B1_ receptor by binding to its promoter region.

### Notch1 signalling regulated MIP through GABA_B1_ receptor

To further test whether Notch1 signalling regulated MIP through the GABA_B1_ receptor, we co-regulated Notch1 and GABA_B_ receptor activity in the mPFC of mice and examined the behaviours of MIP (Fig. 5A). The treatment of phaclofen, a GABA_B_ receptor antagonist, abolished the suppression of METH sensitization induced by syn-NICD-shRNA (Fig. 5B). Moreover, a GABA_B_ receptor agonist baclofen completely normalized the enhancement of METH sensitization by syn-NICD-OE (Fig. 5D). Meanwhile, the effect of NICD expression changes on the time that mice spent in the centre area of the open field was also reversed by phaclofen and baclofen (Fig. 5C, 5E). Based on no expression changes of GABA_B2_ receptor in syn-NICD-shRNA or syn-NICD-OE group (Fig. S4C, S4H), Notch1 signalling may regulate MIP through GABA_B1_ receptors.

**Figure 5.**
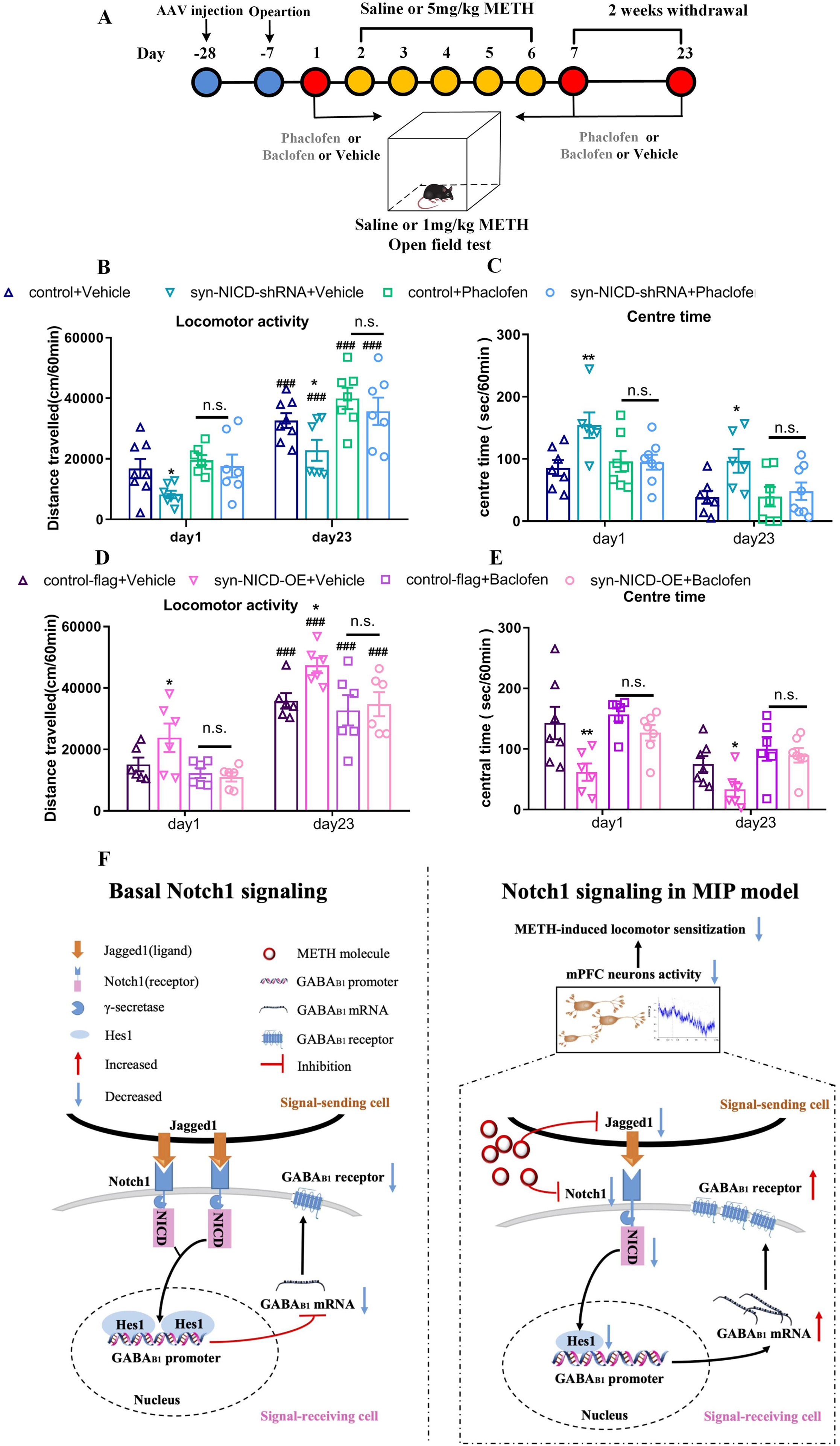
GABA_B_ receptor antagonist (phaclofen) or agonist (baclofen) reversed the effect of Notch1 signalling in METH-induced sensitization. 5A. Timeline of phaclofen or baclofen pre-treatment for the syn-NICD-shRNA or syn-NICD-OE METH group in the locomotor activity test. **5B-5C.** Phaclofen pre-treatment reversed the significant decrease in locomotion and increase in centre time evoked by inhibition of NICD in the mPFC on day 1 and day 23. **5B.** Mixed-design ANOVA followed by post hoc multiple LSD test showed significant main effects of phaclofen [F _(1, 23)_ = 17.61, *P <* 0.001]; AAV [F _(1, 23)_ = 5.10, *P <* 0.05]; and AAV×METH [F _(1, 23)_ = 4.06, *P <* 0.05] on locomotor activity. **5C.** The significant main effect of phaclofen [F _(1, 24)_ = 8.52, *P <* 0.01]; AAV [F _(1, 24)_ = 4.84, *P <* 0.05]; and AAV×METH [F _(1, 24)_ = 7.61, *P <* 0.05] on centre time. **5D-5E.** Baclofen pre-treatment reversed the significant increase in locomotion and decrease in centre time evoked by overexpression of NICD in the mPFC on day 1 and day 23. **5D.** Mixed-design ANOVA with LSD post hoc multiple comparison was performed. There were significant main effects of baclofen [F _(1, 20)_ = 11.31, *P <* 0.01]; AAV [F _(1, 20)_ = 5.11, *P <* 0.05]; and AAV×METH [F _(1, 20)_ = 4.34, *P <* 0.05] on locomotor activity. **5E.** There were significant main effect of baclofen [F _(1, 22)_ = 11.46, *P <* 0.01]; AAV [F _(1, 22)_ = 11.12, *P <* 0.01]; but not the AAV×METH [F _(1, 22)_ = 2.86, *P >* 0.05] on centre time. \**P <* 0.05, \*\**P <* 0.01 vs. the vehicle control group. ###*P* < 0.001 vs. the same group on day1. n.s. means no significant changes. The results are expressed as the mean ± SEM, n = 6-8. **5F.** Working model of Notch1 signalling pathway–mediated regulation of the GABA_B1_ receptor as a protective factor against METH-induced behaviour sensitization. We show that the baseline levels of Notch1 signalling in the mPFC as a consequence of ligand (Jagged1) and receptor (Notch 1) interaction maintain the status quo of the baseline state (left). In the MIP model, Notch1 signalling downregulation in the mPFC results in an increase in the GABA_B1_ receptor, thereby creating an environment that attenuates mPFC neural activity and METH-induced locomotor sensitization.

## Discussion

A mechanistic understanding of how MIP occurs and develops in terms of neuronal molecular events is currently lacking. Our approach to addressing this process is a unique intersection of the study of MIP, Notch1 signalling and GABA receptors. We found that regulating the expression of Notch1 in the mPFC affects MIP by influencing GABA_B1_ receptor expression *via* Hes1-dependent Notch1 signalling mechanisms. To the best of our knowledge, these data provide the first evidence for a potential link between dysregulation of Notch1 signalling in the mPFC and the pathogenesis of MIP.

Some previous studies have reported that the indistinguishable symptoms of MIP and SCZ might share similar risk factors and molecular mechanisms^3, 13^. However, we observed specific and significant decreases in Notch1 signalling components in the mPFC of MIP mice. In contrast, neither animal models of SCZ nor postmortem data from SCZ patients have shown significant changes in Notch1 signalling in the mPFC. These results suggest that MIP may have a specific mechanism that is different from that of SCZ. Besides, synaptic dysfunctions in SCZ and MIP models are distinct; in the MK-801-treated animal model of SCZ, hyperactivity is caused by NMDAR hypofunction, whereas MIP is mainly caused by the overflow of dopamine in synapses^46^. It has been proved that D2 type dopamine receptor (D2R) agonist ropinirole can affect the Notch1 mRNA expression^47^. Hence, it is reasonable to assume that the reduction in Notch1 signalling may be closely related to excessive dopaminergic activity but not NMDA receptor hypofunction. Potential interactions between the Notch1 signalling pathway and dopamine system in the pathogenesis of MIP warrant further study.

We further demonstrate that the downregulation of neural Notch1 in the mPFC attenuates MIP-related behaviours and that Notch1 overexpression in neurons aggravates these MIP-related behaviours (including sensitization, anxiety-like behaviours, depression-like behaviours and cognitive impairment). Consistent with our results, the administration of the DAPT in the mPFC alleviated cognitive deficits in a rat model of autism^48^. NaHS caused remission from the depression- and anxiety-like behaviours induced by type 1 diabetes mellitus by decreasing Notch1 signalling^49^. All of these results suggested that Notch1 signalling in the mPFC may play an important role in MIP and that the reduction in Notch1 signalling could be a protective factor against MIP.

Reduced activity of the GABAergic inhibitory network was one of the key factors in the occurrence of MIP^2, 50^. It has been reported that the activity of the GABAergic system in the mPFC decreased in METH-sensitized mice^27, 51^. Our results further confirmed the significant downregulation of GABA receptors and transporters in the mPFC of MIP mice. Interestingly, through a variety of GABA receptors and transporters expression analyses, we found that Notch1 signalling could negatively regulate GABA_B1_ receptor expression. Furthermore, our results revealed that inhibition of Hes1, a transcriptional repressor of Notch1 signalling, could increase GABA_B1_ receptor expression. Indeed, using ChIP-qPCR assays, we obtained evidence that the downstream transcriptional repressor Hes1 could directly bind to the GABA_B1_ receptor promoters.

GABA_B1_, a subunit of GABA_B_ receptors, is responsible for G-protein coupling and modulates potassium channels and high-voltage calcium channels^52^. Activation of the GABA_B_ receptor could reduce neuronal excitability^53^ and exert an antipsychotic-like action in SCZ^54^. In addition, increasing GABA_B_ receptor activity in the PFC could decrease psychostimulant-induced dopamine levels in the prefrontal cortex^55^ and ameliorate METH-induced locomotor activity^56^ and cognitive deficits^57^. In accordance with these studies, increased GABA_B1_ receptor expression in the mPFC induced by Notch1 signalling suppressed METH-induced locomotor hyperactivity, while decreased GABA_B1_ receptor expression augmented the response to METH-induced locomotor activity. Furthermore, *in vivo* fiber photometry data suggest that the Notch1-shRNA-induced increase in GABA_B1_ receptor may cause the inhibition of mPFC neurons. More notably, pretreatment with a GABA_B_ receptor antagonist or agonist dramatically reversed the Notch1-induced changes in METH sensitization. Given the above, our results suggested that a METH-induced decrease in the expression of Notch1 in the mPFC could increase GABA_B1_ receptor expression and consequently inhibit mPFC activity, resulting in an antipsychotic effect on the MIP mouse model (Fig. 5F). Although both GABA_B1_ receptor and Notch1 signalling were significantly reduced in the mPFC of MIP mice, this phenomenon may be resulting from direct METH influence or other transcription regulators. For example, glutamate can lead to rapid down-regulation of GABA_B_ receptors via lysosomal degradation in cortical neurons^58^. The higher levels of extracellular glutamate in mPFC of METH-acute^59^ and METH-sensitized mice^60^ may directly result in the downregulation of GABA_B1_ receptor rather than Notch1 signalling.

A few interesting questions remain open following this study. First, the type of neuron on which the Notch1-Hes1-GABA_B1_ pathway acts in the mPFC is still unknown. According to our results and previous studies^61^, upregulation of the GABA_B1_ receptor would lead to reduced mPFC activity. Thus, expression changes are most likely to occur in glutamatergic pyramidal neurons. Further investigations should be carried out to test this speculation. Second, the neuronal activity could be detected more precisely through electrophysiology, which could determine whether GABA_B1_ expression changes in the presynaptic or postsynaptic cells. Finally, it would be worthwhile to identify the brain regions to which the activated mPFC neurons project in MIP.

In conclusion, our study presents the first data to demonstrate reduced the expression of the Notch1 signalling pathway in the mPFC of a MIP mouse model. More importantly, the decreased Notch1 signalling in the mPFC of mice was capable of attenuating MIP-related behaviours. These effects can occur through increased GABA_B1_ receptor expression as a result of Notch1-Hes1 signalling, which could inhibit mPFC neuronal activity (Fig. 5F). Our work proposes an important associations between Notch1 signalling and MIP-related neural plasticity. These findings will provide mechanistic insights into understanding of MIP and, ultimately, facilitate the development of new therapies for MIP.

## Acknowledgements

This research was supported by grants from the National Natural Science Foundation of China (No. 82171879, No. 81772034, No. 81772033, No. 82171873, No. 81922024); Science, Technology and Innovation Commission of Shenzhen Municipality (RCJC20200714114556103).

## Author contributions

TN, LZ and SW conducted experiments and analyses. TC, FG and LZ designed experiments and provided materials. TN drafted the manuscript. LZ, FG, WZ, YX, YZ, DM and HW provided critical revisions of the manuscript. All of the authors critically reviewed the content and approved the final version of the manuscript for publication.

## Supplementary information is available at MP’s website

