## supplement figure 1-4 for "Medial prefrontal cortex Notch1 signalling mediates methamphetamine-induced psychosis via Hes1-dependent suppression of GABA_B1_ receptor expression"

### Supplement methods

#### Behavioural Testing

##### Open filed test

Behavioural effects of the MK801 treatments were quantified using the open field test. Individual mice were placed in the metal test chambers (43 × 43 × 43 cm) and analysed the day after last MK-801 injection using a smart 2.5 video tracking system. The total distances moved throughout the area as well as the time spent in the centre zone were recorded for 10 min.

##### Y-maze spontaneous alternation

The Y-maze was a Y-shaped apparatus with three arms, each 30 cm long and 6 cm wide with walls 15 cm high. The arms were at a 120 ° angle from each other. Mice were randomly placed at the end of one of the 3 arms to avoid placement bias and allowed to freely explore the Y-maze for 5 minutes. A correct alternation was defined by a trio of arm entries in which the animal accessed all three arms in sequence without repetition (ABC, BCA, and CAB). Alternation triplet = [(correct alternations)/(total arm entries − 2)] × 100%[1].

##### Novel object recognition task (NOR)

The procedure for this test follows Antunes et al[2].Animals were individually habituated to an empty box (30 × 45 × 16 cm high) for 10 min on two consecutive days. During the training phase, two identical objects (A and A) were placed in the middle area of the chamber. Each animal was then placed in the box and allowed to freely explore the area for 5 min. This trial was repeated three times at 15 min intervals. Three hours after the training phase, one object (A or B randomly) was substituted for a novel object (C), and exploratory behaviour was again evaluated for a total of 5 min in test 1. For the last test, object C was replaced by another novel object (D). During the trials, the open field box and all objects were thoroughly cleansed using 70% ethanol between sessions to prevent odour recognition. Exploration of an object was characterized as sniffing it at a distance of less than 2 cm, and/or touching it with the nose. Smart 3.0 software was used to record the exploration time. Discrimination of visual novelty was assessed by a recognition index, defined as: (new object exploration–old object exploration)/ (total time spent exploring both objects).

##### Elevated plus maze (EPM)

The EPM consisted of a plus-shaped platform with two open (33 × 6 cm) and two closed arms extending from a 6 × 6 cm central area; the platform was elevated 50 cm off the ground. During the 5-min trial, the time spent in the open and closed arms and the number of entries into arms were measured[3] by Smart 3.0 software.

**Forced swimming test (FST)**

The FST is performed to assess despair behavior[4]. Each mouse was placed in a 10-litre transparent plastic cylinder that was filled with tap water at 25 ± 1°C to a depth of 30 cm; the mouse was then left to move freely in the water for 6 min. Immobility was defined as a lack of movements except those necessary to prevent the animal from drowning. The percent immobility was calculated as the percent of immobile behaviour over the test duration. Moreover, the immobility duration was recorded and analysed during the last 5 of the 6 min using Smart 3.0 software.

**Tail suspension test (TST)**

The TST was performed in accordance with previously described methods[4]. Briefly, the mice were suspended with adhesive tape from a hook 50 cm above soft bedding material in a chamber that was both acoustically and visually isolated. The hook was placed approximately 1 cm from the tip of the tail. The mice were suspended for 6 min, and the immobility duration was recorded and analysed during the last 5 min of the 6 min test using Smart 3.0 software.

**Supplement Table 1**. Summary of forward and reverse primer sequences in quantitative real-time PCR

| Gene | Forward primer sequence (5′-3′) | Reverse primer sequence (5′-3′) |
|---|---|---|
| Notch1 | TCAGGGTGTCTTCCAGATCC | CAGCATCCACATTGTTCACC |
| Jagged1 | TGACATGGATAAACACCAGCA | GCAGCCCACTGTCTGCTATAC |
| RBP-J | CCAATTTCAGGCCACTCCA | TCTACATCCCCAAACCACACTC |
| Hes1 | GCAGACATTCTGGAAATGACTGTGA | GAGTGCGCACCTCGGTGTTA |
| GAT1 | TAACAACAACAGCCCATCCA | GGAGTAACCCTGCTCCATGA |
| GAT3 | CCTCCATGATCTGCATTCCT | CCAAATACCCCCTTTCGTCT |
| GABAAα3 | CAAGAACCTGGGGACTTTCTCAA | AGCCGATCCAAGATTCTAGTGAA |
| GABAA β1 | GGTTTGTTGTGCACACAGCTCC | ATGCTGGCGACATCGATCCGC |
| GABAB1 | ACGTCACCTCGGAAGGTTG | CACAGGCAGGAAATTGATGGC |
| GABAB2 | CCTGGTCATCATCTTCTGTAGCA | AACTGGAATCGCCTGTTCTGA |
| Gapdh | TGTGTCCGTCGTGGATCTGA | TTGCTGTTGAAGTCGCAGGAG |
| Primer1(ChIP) | AGCCTACTGCTGGACTAACGA | AGGGAGAATCAAGGGTCAAAA |
| Primer2(ChIP) | ATACCTGCTTTCCACCCAC | CTCCTGCATTGCGATTGTC |
| Primer3(ChIP) | TTAGTTCGTGGTTTAGGAGTC | GTGGAAAGCAGGTATTATGAG |
| Gapdh | TGTGTCCGTCGTGGATCTGA | TTGCTGTTGAAGTCGCAGGAG |

The relative expressions of these genes were normalized to the Gapdh. The melt temperature (Tm) was kept between 55 ℃ and 65 ℃.

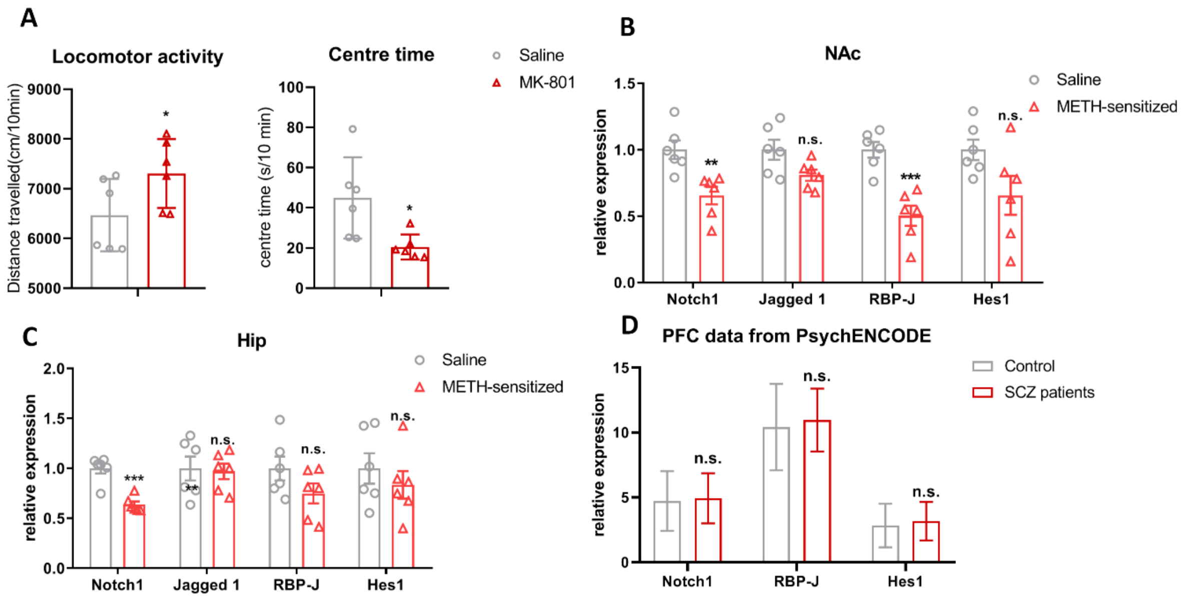


**Supplement figure 1. The expression changes of Notch1 signalling.**

**1A.**The effect of repeated MK-801 treatment in the open filed test. Repeat MK-801 treatment significantly increased locomotor activity ($t_{10} = -2.30$, $P < 0.05$) and decreased the time spend in centre area ($t_{10} = 2.82$, $P < 0.05$). **1B.** mRNA level of the Notch1 signal pathway in the NAc of METH sensitized mice. There was significant downregulation of Notch1 ($t_{10} = 3.61$, $P < 0.01$) and RBP-J ($t_{10} = 5.15$, $P < 0.001$) but not Jagged1 ($t_{10} = 2.22$, $P > 0.05$) or Hes1 ($t_{10} = 2.08$, $P > 0.05$) by student's t test. **1C.** mRNA level of the Notch1 signalling in the Hip of METH-sensitized mice. The Notch1 receptor was significantly reduced ($t_{10} = 5.99$, $P < 0.001$). However, there were no significant changes of Jagged1 ($t_{10} = 0.21$, $P > 0.05$), RBP-J ($t_{10} = 1.61$, $P > 0.05$) or Hes1 ($t_{10} = 0.80$, $P > 0.05$) by student's t test. **1D.** mRNA-sequencing data from the PsychENCODE consortium including 256 healthy people and 95 schizophrenia patients' prefrontal cortex data. There were no significant changes in Notch1 ($t_{349} = -0.80$, $P > 0.05$), RBP-J ($t_{349} = -1.46$, $P > 0.05$) or Hes1 ($t_{349} = -1.67$, $P > 0.05$) between schizophrenia patients and controls by student's t test. *$P < 0.05$, **$P < 0.01$, ***$P < 0.001$ vs. saline group. n.s. means no significant changes . Data were presented as mean ± S.E.M, n = 6.

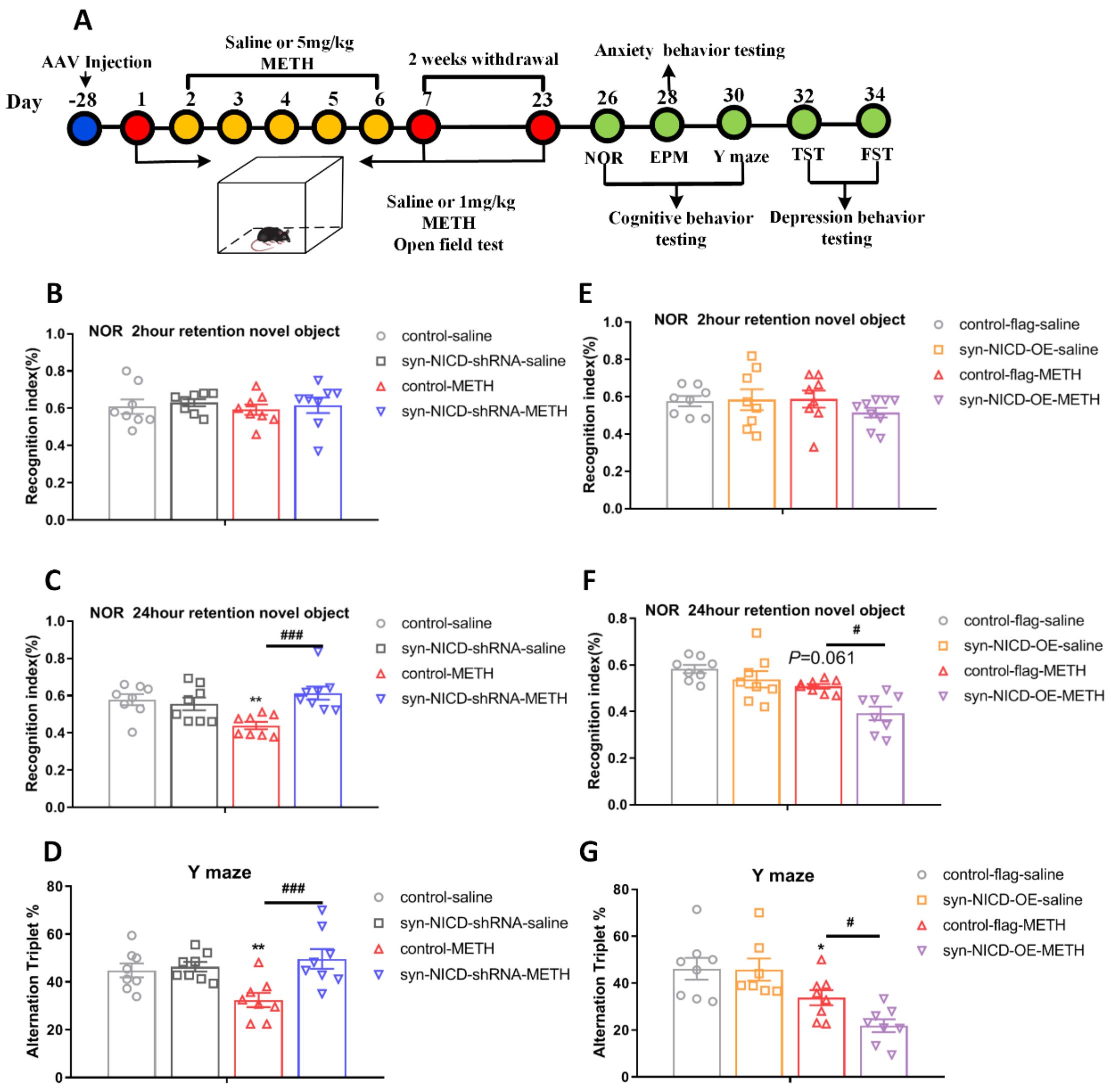


**Supplement figure 2. Manipulating Notch1 expression affects METH-induced psychosis (NOR and Y-maze).**

**2A.** Timeline of MIP relevant behavioural test. These behavioural tests were carried out within 2 weeks after the METH challenge injection. **2B-2C.** Inhibition of Notch1 signalling reversed the decrease in the recognition index (%) after 24 hours of training but not after 2 hours. **2C.** Two-way ANOVA with LSD post hoc multiple comparison revealed significant main effect of AAV [$F_{(1, 28)} = 6.06$, $P < 0.05$] and METH×AAV [$F_{(1, 28)} = 10.48$, $P < 0.01$], but not the METH ($P > 0.05$). **2D.** Downregulation of Notch1 signalling reversed the reduction in alternation triplet in the Y-maze [main effect of AAV, $F_{(1, 28)} = 8.99$, $P < 0.01$; METH, $F_{(1, 28)} = 2.14$, $P > 0.05$; AAV×METH, $F_{(1, 28)} = 6.27$, $P < 0.05$] **2E-2F.** Overexpression of Notch1 signalling enhanced the decreased recognition index (%) after 24 hours of training but not after two hours. **2F.** Two-way

ANOVA with LSD post hoc multiple comparison revealed significant main effect of AAV [$F_{(1, 28)} = 10.11$, $P < 0.01$] and METH [$F_{(1, 28)} = 19.17$, $P < 0.001$], but not the AAV×METH ($P > 0.05$). **2G.** Overexpression of Notch1 signalling aggravated the reduction in alternation triplet in the Y-maze [main effect of AAV, $F_{(1, 27)} = 2.53$, $P > 0.05$; METH, $F_{(1, 27)} = 21.84$, $P < 0.001$; AAV×METH, $F_{(1, 27)} = 2.24$, $P > 0.05$].*$P < 0.05$, **$P < 0.01$ vs. saline control group. #$P < 0.05$, ###$P < 0.001$ control-METH vs. syn-NICD-OE-METH or syn-NICD-shRNA-METH group. Data were presented as mean ± S.E.M, n = 7-8.

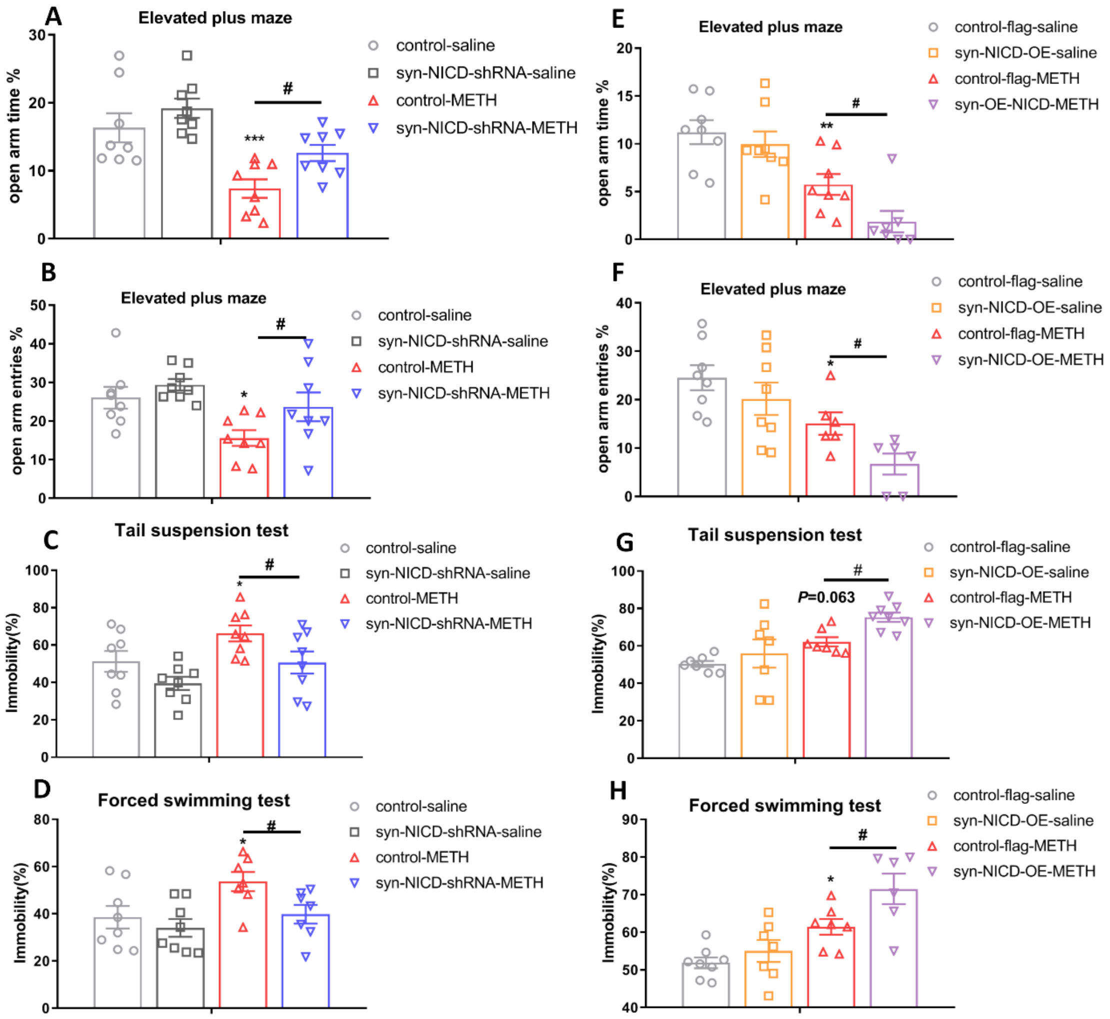


**Supplement figure 3. Manipulating Notch1 expression affects METH-induced psychosis (EPM, TST and FST).**

**3A-3B, 3E-3F.** MIP mice spent a significantly lower proportion of time in the open arms and made a lower proportion of entries into the open arms than the control saline group, indicating a higher level of anxiety in MIP mice. **3A-3B.** Syn-NICD-shRNA-METH group of mice reversed the decrease time that spent in the open arms [**3A**, main effect of AAV, $F_{(1, 28)}$=6.60, $P < 0.05$; METH, $F_{(1, 28)} = 21.19$, $P < 0.001$; AAV ×METH, $P$>0.05] and the number of entries into the open arms [**3B**, main effect of AAV, $F_{(1, 28)} = 4.61$, $P < 0.05$; METH, $F_{(1, 28)} = 9.25$, $P < 0.01$; AAV ×METH, $P > 0.05$ ]. **3E-3F.** Syn-NICD-OE-METH group of mice aggravated the decreased time that spent in the open arms [**3E**, main effect of AAV, $F_{(1, 27)} = 4.45$, $P < 0.05$; METH ,$F_{(1, 27)} = 31.03$, $P < 0.001$; AAV ×METH, $P > 0.05$] and the number of entries in the open arms [**3F**, main effect of AAV, $F_{(1, 24)} = 5.15$, $P < 0.05$; METH, $F_{(1, 24)} = 16.67$, $P < 0.001$; AAV ×METH, $P > 0.05$]. **3C-3D, 3G-3H.** METH-treated mice had greater percentages of immobility time in the TST and FST than control mice. The syn-NICD-shRNA-METH group of mice reversed the increased percentage of immobility time in the TST and FST. [**3C**, main effect of AAV, $F_{(1, 28)} = 7.65$, $P < 0.05$; METH, $F_{(1, 28)} = 7.00$, $P < 0.05$; AAV ×METH, $P > 0.05$; **3D**. main effect of AAV, $F_{(1, 25)} = 5.14$, $P < 0.05$; METH, $F_{(1, 25)} = 14.21$, $P<0.01$; AAV ×METH, $P > 0.05$]. **3G-3H.** The syn-NICD-OE-METH group of mice exhibited an increased percentage of immobility time in the TST and FST [**3G**, main effect of AAV, $F_{(1, 26)} = 4.92$, $P < 0.05$; METH, $F_{(1, 26)} = 4.19$, $P = 0.051$; AAV ×METH, $P > 0.05$; **3H.** main effect of AAV, $F_{(1, 24)} = 6.31$, $P < 0.05$; METH, $F_{(1, 24)} = 24.52$, $P < 0.001$; AAV ×METH, $P > 0.05$]. *$P < 0.05$, **$P < 0.01$ ***$P < 0.001$ vs. saline control group. #$P < 0.05$ vs. the METH control group. Data were presented as mean ± S.E.M, n = 7-8.

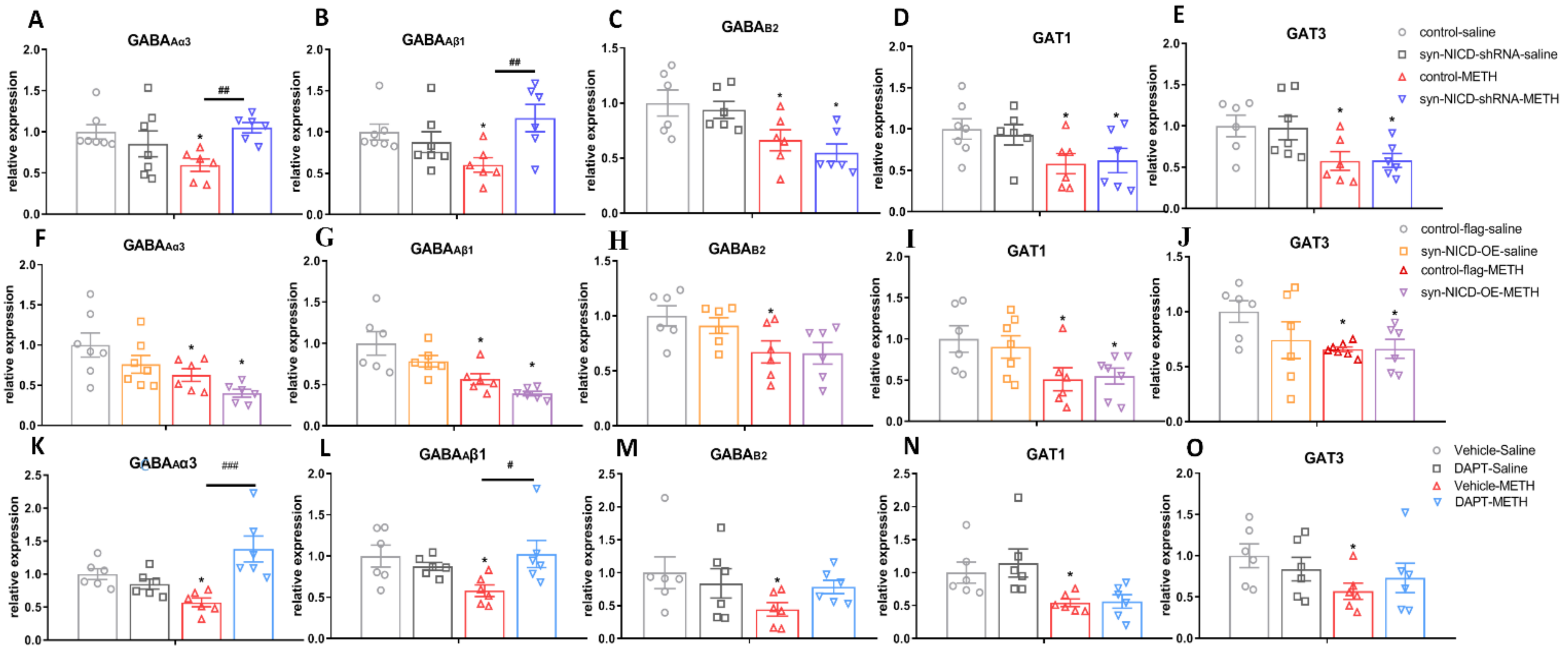


**Supplement figure 4. Changes in the expression of GABA receptors and transporters in the mPFC with the manipulation of mPFC Noch1 signalling in MIP mice.**

**4A-4E.** Changes in the expression of the GABAergic system in the syn-NICD-shRNA-saline and METH groups of mice. Two-way ANOVA followed LSD multiple comparisons. METH caused a significant decrease in $GABA_{A\alpha3}$, $GABA_{A\beta1}$ $GABA_{B2}$, GAT1 and GAT3. The syn-NICD-shRNA-METH group only exhibited upregulation of $GABA_{A\alpha3}$ and $GABA_{A\beta1}$ receptor expression compared with the control METH group. **4A.**

$GABA_{A\alpha3}$, the main effect of AAV, $F_{(1, 22)} = 2.08$, $P > 0.05$; METH, $F_{(1, 22)} = 0.94$, $P > 0.05$; interaction, $F_{(1, 22)} = 7.83$, $P < 0.05$. **4B.** $GABA_{A\beta1}$, the main effect of AAV, $F_{(1, 22)} = 3.31$, $P > 0.05$; METH, $F_{(1, 22)} = 0.18$, $P > 0.05$; interaction $F_{(1, 22)} = 8.00$, $P < 0.05$. **4C.** $GABA_{B2}$, the main effect of AAV, $F_{(1, 20)} = 0.87$, $P > 0.05$; METH, $F_{(1, 20)} = 14.97$, $P < 0.01$; interaction, $F_{(1, 20)} = 0.08$, $P > 0.05$. **4D.** GAT1, the main effect of AAV, $F_{(1, 21)} = 0.02$, $P > 0.05$; METH, $F_{(1, 21)} = 7.88$, $P < 0.05$; interaction, $F_{(1, 21)} = 0.17$, $P > 0.05$. **4E.** GAT3, the main effect of AAV, $F_{(1, 23)} = 0.02$, $P > 0.05$; METH, $F_{(1, 23)} = 11.33$, $P < 0.01$; interaction, $F_{(1, 23)} = 0.004$, $P > 0.05$. **4F-4J.** Changes in the expression of the GABAergic system in the syn-NICD-OE-saline and METH groups of mice. There were no significant changes between syn-NICD-OE-METH and control-flag-METH group on these genes. **4F.** $GABA_{A\alpha3}$, the main effect of AAV, $F_{(1, 22)} = 4.47$, $P < 0.05$; METH, $F_{(1, 22)} =11.07$, $P < 0.05$; interaction, $F_{(1, 22)} = 0.004$, $P > 0.05$. **4G.** $GABA_{A\beta1}$, the main effect of AAV, $F_{(1, 21)} = 6.10$, $P < 0.05$; METH, $F_{(1, 21)} = 21.98$, $P < 0.05$; interaction, $F_{(1, 21)} = 0.46$, $P > 0.05$. **4H.** $GABA_{B2}$, the main effect of AAV, $F_{(1, 20)} = 0.31$, $P > 0.05$; METH, $F_{(1, 20)} = 10.03$, $P < 0.01$; interaction, $F_{(1, 20)} = 0.18$, $P > 0.05$. **4I.** GAT1, the main effect of AAV, $F_{(1, 22)} = 0.05$, $P > 0.05$; METH, $F_{(1, 22)} = 10.01$, $P < 0.01$; interaction, $F_{(1, 22)} = 0.25$, $P > 0.05$. **4J.** GAT3, the main effect of AAV, $F_{(1, 21)} = 1.55$, $P > 0.05$; METH, $F_{(1, 21)} = 4.50$, $P = 0.05$; interaction, $F_{(1, 23)} = 1.65$, $P > 0.05$. **4K-4O.** The GABAergic system expression changes following administration of DAPT in saline and METH group of mice. The DAPT-METH only show upregulation of $GABA_{A\alpha3}$ and $GABA_{A\beta1}$ receptor expression compared with the Vehicle-METH group. **4K.** $GABA_{A\alpha3}$, the main effect of DAPT, $F_{(1, 20)} = 7.91$, $P < 0.05$; METH, $F_{(1,20)} = 0.20$, $P > 0.05$; interaction, $F_{(1, 20)} = 16.72$, $P < 0.001$; **4L**. $GABA_{A\beta1}$, the main effect of DAPT, $F_{(1, 20)} = 1.36$, $P > 0.05$; METH, $F_{(1, 20)} = 1.95$, $P > 0.05$; interaction, $F_{(1, 20)} = 6.22$, $P < 0.05$; **4M.** $GABA_{B2}$, the main effect of DAPT, $F_{(1, 20)} = 0.25$, $P > 0.05$; METH, $F_{(1, 20)} = 2.88$, $P > 0.05$; interaction, $F_{(1, 20)} = 1.95$, $P > 0.05$. **4N.** For GAT1, the main effect of DAPT, $F_{(1, 23)} = 0.32$, $P > 0.05$; METH, $F_{(1, 20)} = 12.89$, $P < 0.01$; interaction, $F_{(1, 23)} = 0.19$, $P > 0.05$. **4O.** For GAT3, the main effect of DAPT, $F_{(1, 20)} = 0.001$, $P > 0.05$, METH, $F_{(1, 20)} = 3.48$, $P > 0.05$, interaction, $F_{(1, 23)} = 1.26$, $P > 0.05$. $^*P < 0.05$, compared to the paired saline group. $\#P < 0.05$, $\#\# P < 0.01$, $\#\#\#P < 0.001$ compared to the paired METH group. Data were presented as mean ± S.E.M, n = 6-7.
